# The march of the Common Green Iguana (*Iguana iguana*): early establishment in Singapore and Thailand is facilitated by the pet trade and recreational parks

**DOI:** 10.1101/2020.02.04.933598

**Authors:** Matthijs P. van den Burg, Steven M. Van Belleghem, Christina N. De Jesús Villanueva

## Abstract

The popularity of the Common Green Iguana (*Iguana iguana*) as a pet has contributed to its global occurrence as an invasive alien species. Early detection and control of invasive alien *I. iguana* populations is necessary to prevent the need for large and financially demanding eradication actions. Here, we collated information from digital footage and interviews regarding sightings of free roaming *I. iguana* specimens in Singapore and Thailand, and present evidence of early-stage invasions and establishment. Using species distribution modeling, we find that large parts of Thailand and neighboring countries have suitable habitat, which could facilitate the expansion of these alien populations if left uncontrolled. Additionally, we report singular *I. iguana* sightings in Hong Kong and Peninsular Malaysia. We call for awareness of alien *I. iguana* in the Philippines due to the high number of pet iguanas and reported CITES importations as well as the availability of suitable habitat throughout the archipelago. Further, we identify *I. iguana* presence to be facilitated by the release of pet-traded specimens and uncontrolled exhibition practices in recreational parks. We provide recommendations for implementing monitoring and eradication efforts and strategy recommendations to halt future spread and release.

## Introduction

The Common Green Iguana (*Iguana iguana*, Linnaeus, 1758) is a heavily traded reptilian pet species that has seen its global transport regulated since 1977 (CITES 2019). Despite regulation, the pet trade is an important contributor to the origin of invasive populations (Falcón et al. 2013; Bock et al. 2018; De Jesús Villanueva et al. under review), although other introduction facilitators are known (Censky et al. 1998; van den Burg et al. 2020). *Iguana iguana* occurs throughout Central and South America as well as on several Caribbean islands (Stephen et al. 2013; Bock et al. 2018). Invasions of *I. iguana* are mainly restricted to Caribbean islands, with only a small set of islands known from outside this region: Canary Islands (Spain), Fiji, Hawaii (USA), Japan and Taiwan (Bock et al. 2018; Lee et al. 2019). Importantly, established populations of this invasive alien species (IAS) have never been successfully eradicated (Lee et al. 2019; Rivera-Milán and Haakonsson in press).

The rapid growth of *I. iguana* populations can pose a threat to the economy of invaded territories, through infrastructure damage and agricultural loss, as well as threats to biodiversity (Bock et al. 2018). As invasive populations can originate from multiple genetic lineages (Stephen et al. 2013; van den Burg et al. 2018a; De Jesús Villanueva et al. under review) high genetic diversity might increase adaptive variation to persevere in non-native environments (Kolbe et al. 2004). In Puerto Rico, and presumably other invasive areas, *I. iguana* population growth may be further aided by a low number of predators and direct competitors (López-Torres et al. 2011). Furthermore, invasive *I. iguana* populations are a threat to native species through direct competition and hybridization (Vuillaume et al. 2015; Moss et al. 2018; van den Burg et al. 2018b; van Wagensveld and van den Burg 2018). On Grand Cayman, an island of 196 km^2^, the present invasive *I. iguana* population was estimated to reach between 0.9–1.9 million individuals in early 2018 before the start of a government-financed culling program (Rivera-Milán and Haakonsson in press). During this program’s initial year, through collective work of 500 hunters, over 960,000 iguanas were removed against the expense of 5.7 million USD (Rivera-Milán and Haakonsson in press). To prevent further spread of this species and the need for multimillion-dollar eradication programs, rapid management action is key during the initial introduction and establishment phase (Davis 2009; Blackburn et al. 2011).

In southeast Asia, Thailand and Singapore are considered important hubs in the international wildlife trade (Nijman 2010), with herpetofaunal survey reports having identified notable discrepancies among CITES import data, and the presence of several non-native species in the wild in these countries (Nijman and Shepherd 2007, 2010, 2011; Yeo and Chia 2010; Poole and Shepherd 2016). For Thailand, Nijman and Shepherd (2011) found that over 150 distinct species of amphibians and reptiles had CITES import records between 1990–2007. In contrast, only two reptiles (*Trachemys scripta elegans* and *I. iguana*) were recorded in Thailand during a 2011–2012 inventory of wild alien vertebrate fauna (Boonkuaw et al. 2014). For *I. iguana*, Boonkuaw et al. (2014) reported three sightings of single specimens in Thailand, all of which had disappeared before the report’s publication. Singapore is mainly regarded as an import and re-export country (Nijman et al. 2012) and was the main regional reptile importer between 1998–2007 (Nijman 2010). In contrast to Thailand, over 20 alien reptile species were present in Singapore in 2010 (Yeo and Chia 2010). Of these species, only four were considered established (*Calotes versicolor, Pareas margaritophorus, Siebenrockiella crassicollis* and *Xenochrophis vittatus*); with *I. iguana* known from only a few scant recordings (Chua 2007; Yeo and Chia 2010). However, despite previous reports of few *I. iguana* specimens (Yeo and Chia 2010; Boonkuaw et al. 2014), recent reports of free-roaming *I. iguana* on social media platforms (e.g. FaceBook and Instagram) and on iNaturalist have sparked our concern over the potential for established and growing invasive alien populations to be present in Singapore and Thailand. Here, we 1) summarize the current extent of *Iguana iguana* in Singapore and Thailand, 2) identify locations with signs of early establishment and its facilitators, 3) predict the potential range in the absence of future eradication action, and 4) provide recommendations to prevent future releases and mitigate negative ecological effects.

## Methods

### Observations

We traced the presence of alien *I. iguana* in Singapore and Thailand through verifiable sightings using photographs and videos from social media (FaceBook and Instagram), Internet websites (www.flickr.com,www.istockphoto.com, www.shutterstock.com) and iNaturalist. We contacted and interviewed owners of online images whose content suggested the presence of free-roaming *I. iguana* to acquire more details including the location where the picture was taken, date, presence of free roaming *I. iguana*, presence of different *I. iguana* life stages and number of individuals present. Through snowball sampling (where interviewees had more footage or contacts with similar footage) we were able to acquire additional verifiable sightings not publically available. When images were taken within zoos or recreational parks (hereafter recreational parks) we asked whether *I. iguana* were kept in holding cages or were freely roaming. In the latter case, we recorded these facilities as potential sources for current and future invasive populations.

### Suitable habitat modeling

To better understand the potential range of *Iguana iguana* in Singapore and Thailand as well as surrounding countries, we used MaxEnt (Philips et al. 2018) as implemented in the *dismo* R package (Hijmans et al. 2017; R Core Team 2019) to build species distribution models (SDM), for current and future climate scenarios. MaxEnt uses environmental and climatic data from species occurrence localities to build a predictive SDM of habitat suitability (HS) for the study species, which runs between 0 (low suitability) and 1 (high suitability). First, we acquired *I. iguana* occurrence data uploaded to the Global Biodiversity Information Facility (GBIF 2019) and iNaturalist. We filtered data records based on georeference inaccuracy (excluded when > 50 km), added records collected during recent fieldwork by the authors (for Colombia and Puerto Rico) and from recent publications on invasive populations (Kraus 2019; Lee et al. 2019), as well as locations in Singapore and Thailand where hatchling or juvenile *I. iguana* have been observed. We removed duplicated locations and obviously erroneous records from locations without *I. iguana* breeding populations (e.g. Galapagos) and with altitude records outside of the species range (> 1000 meters; Stephen et al. 2013). Remaining records were then filtered to retain only one record per 50 km^2^ (Boria et al. 2014) to remove distribution skewness in locality data, leaving a total of 420 records for final model construction. We retained occurrence data from established alien populations arguing that, although invasive areas can hold different climatic compositions compared to native areas, local climatic variables are suitable for successful reproduction and thus help to generalize intraspecific variation within *I. iguana* (Elith et al. 2010; Briscoe Runquist et al. 2019).

The initial SDM model was built using uncorrelated WorldClim 2.0 bioclimatic variables and altitude layer (Fick and Hijmans 2017; at 2.5 arc-minutes resolution). As correlation between variables can lead to overfitting, we implemented the *vifstep* function within the *sdm* package (Naimi and Araújo 2016). This stepwise process identifies the variance inflation factor (VIF), a measure of how much one variable can be explained by the others, for each variable and removes those with high VIF values (here set to 10; Chatterjee and Hadi 2006). After stepwise filtering, 10 variables were retained for the final model with a 2.0–9.1 range of remaining VIF scores: mean diurnal range (BIO2), isothermality (BIO3), mean temperature of wettest quarter (BIO8), mean temperature of driest quarter (BIO9), precipitation of wettest month (BIO13), precipitation of driest month (BIO14), precipitation seasonality (BIO15), precipitation of warmest quarter (BIO18), precipitation of coldest quarter (BIO19), and altitude.

Bioclimatic and occurrence data were next combined to build a presence-only SDM using MaxEnt (Philips et al. 2018) for both the current and a future climate scenario. These models were generated for southeast Asia, as well as the Philippines given the high number of *I. iguana* footage on social media platforms. Predictions of HS for a future climate scenario were run for 2050 using the AC model and rcp45 for greenhouse gas emission (Fick and Hijmans 2017). For both the current and future climate scenario, a set of 10 SDM replications were run with 80% of records used for model training and 20% for model testing. Model performance was evaluated using the area under the curve summary statistic (AUC) with one-tenth of the data; AUC values of ≥ 0.90 indicate high model performance (Swets 1988). The HS absence threshold was determined from the training specificity and sensitivity as implemented in the *dismo* package (Hijmans et al. 2017).

## Results

For Thailand, sighting information was collected from contact with 15 persons and iNaturalist for records since 2016. Sightings originated from a total of 12 locations distributed throughout Thailand (Fig. 1) with a minimum number of total sightings of 97 *I. iguana* (Table A1, Supporting Information). Adult or subadult life stages were observed at all locations, whereas hatchling or juvenile life stages occurred at four locations. The highest number of sightings by different observers was found around Nakhon Ratchasima town (Thai name: □□□□□□□□□□) in Nakhon Ratchasima province. Locations with the highest number of unique *I. iguana* were Nakhon Si Thammarath and Ubonrat, and in the Uthai Thani district. Lastly, at least four recreational parks exhibit non-caged *I. iguana*: Chai Nat Bird Park, Chiang Mai Night Safari, Nakhon Ratchasima Zoo, and Tha Lat Bird Park.

**Fig. 1.**
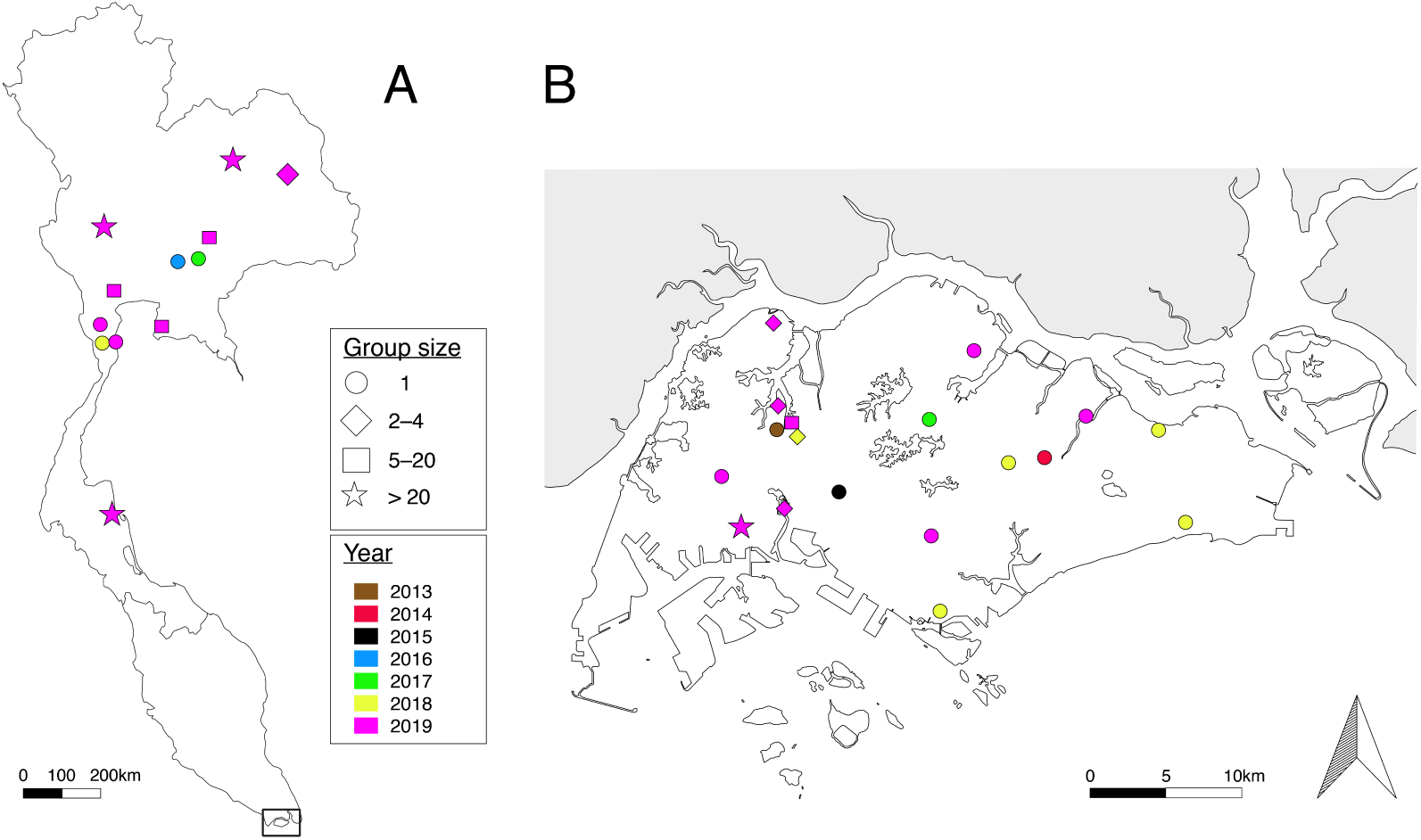
Localities with presence of *Iguana iguana* in (A) Thailand and (B) Singapore showing number of iguanas and year of sighting. Color of the most recent year is displayed per location.

For Singapore, sighting information was collected from iNaturalist records, contact with 10 persons, and from seven unique grey literature publications since 2007 (Chua 2007; Yeo 2014; Ng and Lim 2015; Tay 2015; Khoo 2016; Chew and Low 2017; Yeo 2019). We recorded a minimum of 50 *I. iguana* sightings from 17 locations in Singapore (Fig. 1), of which five locations showed presence of hatchlings and/or juveniles. The highest number of *I. iguana* sightings comes from the area surrounding Jurong hill and Jurong Bird Park as well as around the Warren Golf & Country Club. Recreational parks were not found to exhibit free-roaming *I. iguana* in Singapore.

Besides free-roaming *I. iguana* records from Singapore and Thailand, we recorded two additional locations in southeast Asia. First, one free roaming adult *I. iguana* was observed southeast of Kuala Lumpur, Peninsular Malaysia; this animal was caught in the yard of the observer and was released in an area nearby. The second record is from two free roaming adult iguanas adjacent to the Tsing Tam Reservoirs in Hong Kong.

The SDMs for both current and future climatic conditions suggest the presence of substantial area with *I. iguana* suitable habitat in Thailand and surrounding countries (Fig. 2 and Fig. A1). Suitable habitat is especially present in mainland regions below 15 degrees latitude, except for some areas in Thailand, south and central Peninsular Malaysia and non-coastal regions in Vietnam. Several higher latitude regions have suitable habitat as well including parts of the Gulf of Martaban, and lower elevation areas in Thailand, southern Laos, and central Vietnam. Under a 2050 climate scenario HS in only some areas will substantially change with less areas of lower HS in the Gulf of Martaban, Peninsular Malaysia and Sumatra (Fig. A1, Supporting Information). All *I. iguana* locations reported here below 14 degrees latitude are in high HS areas, with those at higher latitudes in areas with marginal or low HS, except for the Uthai Thani district (Fig. 2). For the Philippines, the current climate model identified suitable habitat in lowland areas on all major islands and marginal habitat or absence of suitable habitat for higher elevations, especially on Mindanao. The median AUC for 10 SDM replications for current climatic conditions was 0.944 (50% interquartile ranges of 0.942–0.946), with a median HS absence threshold of 0.21; and for future climate conditions (2050) 0.948 (50% interquartile ranges of 0.942–0.951) and median HS threshold of 0.28.

**Fig. 2.**
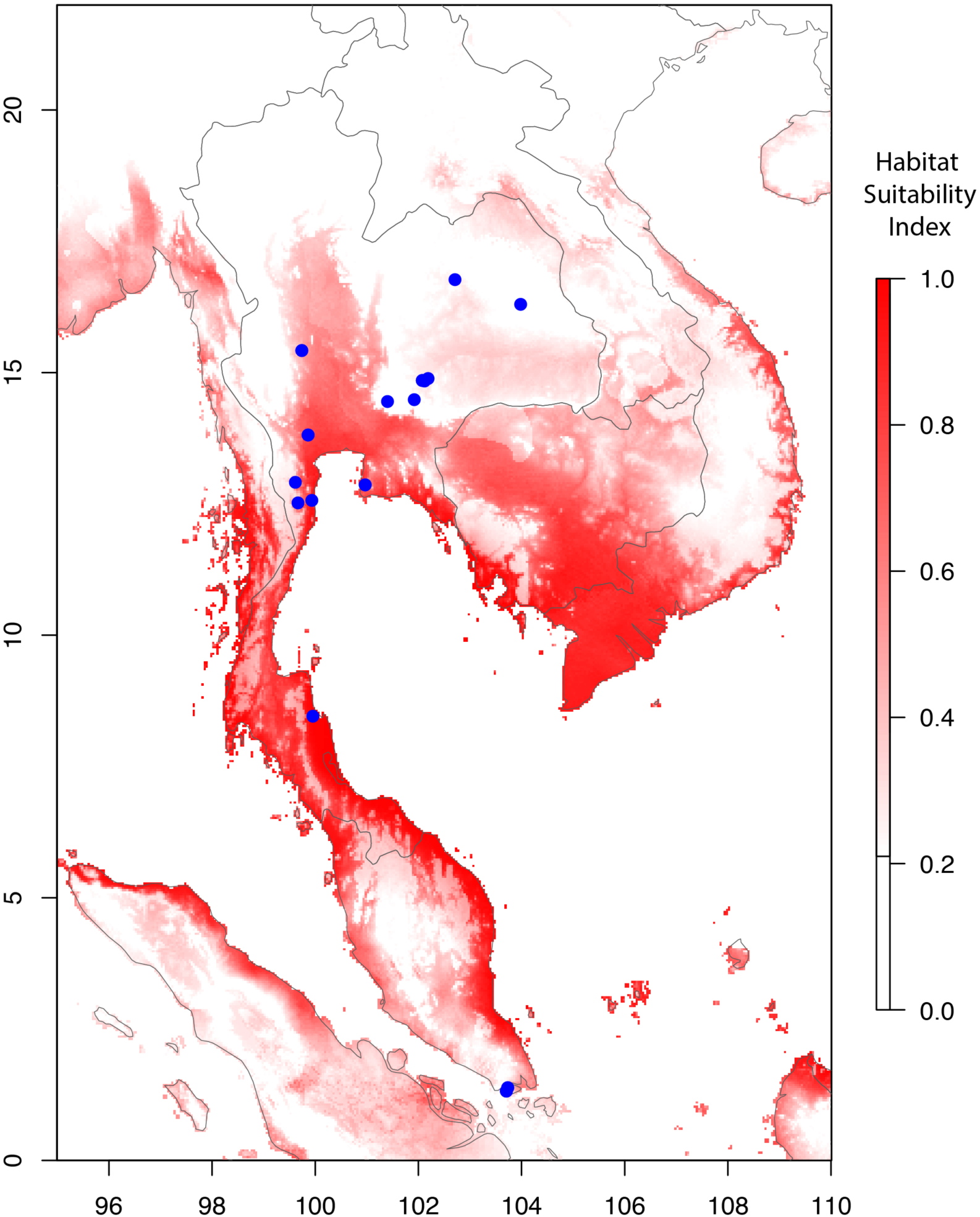
Species distribution models for *Iguana iguana* in Singapore, Thailand and surrounding countries. Models used an absence threshold of 0.17 and habitat suitability ranges from high (dark red) to low (white). Blue dots represent locations with *I. iguana* presence

## Discussion

We report *I. iguana* invasions and early establishment in both Singapore and Thailand with details about population size as well as the current and potential future distribution. In both countries, the presence of hatchling and/or juvenile life stages indicates local recruitment and ongoing establishment of this known invasive alien species (IAS). Importantly, this report describes the second recorded mainland establishment of *I. iguana*, and increases the number of known non-western hemisphere establishments to six; Hawaii, Fiji, Japan, Singapore, Thailand and Taiwan (Mito and Uesugi 2004; Engeman et al. 2011; Falcón et al. 2013; Kraus 2019; Lee et al. 2019); though updates on the status of the Japanese and Hawaiian populations are necessary. Following interviews, we identified the likely source of these invasions to be pet-trade animals that either escaped or were intentionally released by owners or unsecurely exhibited in recreational parks. Besides the release of pet *I. iguana* due to their size, another motive can be fang sheng; the Buddhist act of releasing live animals believed to build spiritual merit (Ng and Lim 2010). As *I. iguana* natively occurs in sub-tropical and tropical biomes identical to those found throughout southeast Asia (Olson et al. 2001), we expect continued spread and growth of these alien populations in the absence of mitigation actions. Indeed, our current and future climate SDMs indicate that large areas in the region have highly suitable *I. iguana* habitat. Additional to over-land dispersal and facilitated by this species’ swim and raft capabilities (Censky et al. 1998; Breuil 1999; van Veen 2011; F. Kraus, pers. comm. 2020), through over-water dispersal a large number of novel locations with suitable habitat could be reached from Singapore (Fig. 2): Peninsular Malaysia, Sumatra, and > 3,000 islands of the Indonesian Riau archipelago. In the absence of removal actions, the persistence and uncontrolled growth of *I. iguana* populations is evident as numerous locations are within highly suitable habitat and recruitment is present (Fig. 2). The case presented here calls for immediate management action as 1) establishment still appears to be in an early phase, 2) eradication expenses are relatively low, 3) current records are likely an underestimate as data were collected opportunistically by non-iguana specialists, 4) a large uncontrolled *I. iguana* population could have far reaching consequences for native biodiversity.

Knowledge on the invasion pathway and source of alien populations is important to prevent repetitive (un)intentional incursions. Although *I. iguana* is native to the American tropics, their importation, captive breeding and escape or release practices could lead to unmanageable population growth in southeast Asia. Between 2000 and 2017, a total of 1,254 live *I. iguana* were imported into Thailand (importer reported quantity; CITES 2019), though currently the number of iguanas present could arguably be higher due to incountry captive breeding practices as well as known import discrepancies in CITES reporting (Nijman and Shepherd 2011). For Singapore, 542 live specimens have been reported by importers pre-2004 (CITES 2019), with no records since, likely due to the country’s illegalization on pet ownership. Several interviewees stated that either they themselves or others (i.e., iguana farms, recreational parks or private residence) have released captive iguanas. This was in addition to deliberate release statements by social media users. Increasing the public awareness of the negative effects from IAS populations may help to discourage future releases.

Uncontrolled or flawed exhibition practices of animals can lead to their escape. In Spain, Fàbregas et al. (2010) identified that enclosure security was not sufficient to prevent escape in 14 % of assessed enclosures, mainly for enclosures holding alien species, which might source IAS populations. For Thailand, based on the presence of free-roaming iguanas close to Thai recreational parks that exhibit non-caged iguanas, these facilities are the most likely sources of iguanas reported from at least two areas: Nakhon Si Thammarath and Nakhon Ratchasima. In Singapore, although we did not acquire information about recreational parks that exhibit free-roaming iguanas, grounds of one park (Jurong Bird Park) are invaded by iguanas. Anecdotal evidence suggests that their presence in the park and surrounding areas originates from the former neighboring reptile recreational park (Jurong Reptile Park). In addition to the risk of escape from parks, free-roaming *I. iguana* within recreational parks are reproductively unhindered, which leads to nesting and successful recruitment. From such nests, small free roaming hatchlings emerge capable of establishing off-ground populations. This poses a particular detection challenge. On top of their small size, the lizards’ green coloration allows them to inconspicuously blend in with the foliage, evading detection for up to a year into their development. In contrast to adults, these hatchlings are harder to catch and track given their small size and capacity to disperse > 200 m within one month after hatching (Knapp and Abarca 2009).

With little information available for Asian *I. iguana* introductions, it is not possible to make straightforward predictions about the ecological impacts of these novel establishments. Data from the western hemisphere demonstrates the possibility that established populations can spread pet-trade diseases to native species (Hellebuyk et al. 2017), competitively displace native species (van den Burg et al. 2018b), and cause erosion and damage to agricultural fields, roads and airplanes (López-Torres et al. 2011). In Thailand we note that invasive populations could have the capacity to displace *Physignathus cocincinus*, the Chinese water dragon (IUCN status Vulnerable; Stuart et al. 2019), given its smaller maximum snout-vent length (SVL) and ecological niche overlap between the species. Namely, *P. cocincinus* is also an arboreal lizard that resides close to streams in east and southeastern Thailand (Stuart et al. 2019). Further, hatchling iguanas will likely be predated by numerous native species given the high regional snake diversity (Chan-Ard et al. 2015), though predators for adult *I. iguana* (which can reach > 45 cm SVL) are few (e.g. large *Python bivittatus* or *Malayopython reticulatus* [Low et al. 2016]). Additionally, *I. iguana* might compete with native species for territory (Chua 2007) and nesting sites (e.g. *Varanus* sp.). However, data on the reproductive cycle of *I. iguana* in Singapore and Thailand is currently lacking, making reproductive period comparisons with native species difficult. Nevertheless, images of mating iguanas as well as a released developed egg suggest mating occurs around November–December. Lastly, high numbers of alien *I. iguana* would presumably cause damage to agricultural fields, gardens and plant nurseries as reported elsewhere (for an overview see Falcón et al. 2013).

For natural areas, over 80% of mangrove plants in a Puerto Rican area were heavily affected by alien *I. iguana* (Carlo and García-Quijano 2008). As mangroves are globally threatened and in decline (Polidoro et al. 2010), also in Singapore (Friess and Webb 2013) despite a recent minor restoration (Lai et al. 2015), alien *I. iguana* are expected to aggravate the decline of Singapore’s mangroves. Recorded *I. iguana* presence in Singapore’s Sungei Buloh Wetland Reserve is therefore worrisome. Future surveys and monitoring are necessary to better understand the ecological and economic impact of *I. iguana* establishment in Singapore and Thailand.

As the global pet trade has facilitated the spread of *I. iguana*, understanding trade dynamics might aid the prediction of future invasions (Robinson et al. 2015; Lockwood et al. 2019). Although the global *I. iguana* trade peaked in the early 2000’s (Stephen et al. 2011), the import of live *I. iguana* to Asia was highest in 2015 and a clear decline is absent (Fig. A2, Supporting Information). We propose that this stable trend is partially due to lower shipping costs and a growing Asian middle class (Kharas 2017). In addition to our report, sightings of released pet iguanas have occurred in Taiwan since 2004 where eradication actions have already removed > 2,000 *I. iguana* individuals (Lee et al. 2019). Similar eradication actions (planned or ongoing) of pet-origin invasive *I. iguana* populations are known from Fiji and Grand Cayman (Kraus 2019; Rivera-Milán and Haakonsson *in press*). Combining these data, we predict the (future) occurrence of additional invasive alien *I. iguana* populations within Asia, specifically in the Philippines, given the apparent high occurrence of pet *I. iguanas* from social media platforms, high number of CITES import records (Sy 2015; CITES 2019), as well as large areas with high habitat suitability (Fig. A3, Supporting Information). Lastly, the records of adult free-roaming iguanas from Peninsular Malaysia and Hong Kong are cause for concern as well. Future assessments should identify whether these are isolated cases or not.

Assertive rapid action is required to halt the spread and to successfully remove alien *I. iguana* populations before they become invasive. To that end we provide several recommendations. First, to mitigate current and future negative effects of these *I. iguana* invasions, we recommend the immediate implementation of survey and questionnaire efforts to better understand the spread of this species in Singapore and Thailand. We recommend both diurnal and nocturnal surveying where diurnal efforts mainly aim to locate adult and sub-adult specimens and nocturnal surveys hatchling and juvenile specimens. Using high-output head torches (e.g. LedLenser H14r.2), small-sized iguanas can be easily located among foliage where they sleep. Secondly, and in parallel, we recommend an outreach campaign to raise public awareness, inform (future) pet owners of all the consequences caused by releasing pet iguanas and promote reporting of *I. iguana* sightings to the relevant (local) authority. Also, information should be provided to new iguana-pet owners upon purchase, addressing the increase in body size, longevity (19.8 years [de Magalhães and Costa 2009]), overall expenses and necessary space. To strengthen this, *I. iguana* retailers should be obligated to exhibit one adult individual (Maceda-Veiga et al. 2019), and we recommend that retailers of exotic pets are obligated to have proven knowledge of this species and of their exhibition standards. Thirdly, we recommend inspections of *I. iguana* captivity circumstances for all facilities where this species is kept; for example, zoos, recreational parks and iguana farms. Iguanas should not be allowed to roam freely, but instead be exhibited in large cages or small-sized open areas enclosed by a deep trench with a smooth metallic vertical surface to prevent climbing; tree branches from within the exhibition area should not extend beyond this trench. Importantly, enclosure or entrapment by water is ineffective given the swimming capacity of *I. iguana*. Lastly, we recommend that releasing a pet *I. iguana* be immediately prohibited by law in Thailand, although despite its prohibition in Singapore the practice continues (Ng and Lim 2010). Pet release prevention could be assisted by pit tagging all imported and in-country bred iguanas to identify registered owners. Besides the above mentioned actions, additional country specific efforts are necessary during this early establishment phase in order to prevent future multi-million dollar eradication efforts.

## Data Accessibility

The R script and data to run SDM models as mentioned in our Methods and Materials are available through GitHub; github.com/StevenVB12/SDM_Iguana.

## Acknowledgements

We are grateful to all contributors for sharing data on *Iguana iguana* sightings. And wish to thank Jason Kolbe for improving an older version of this manuscript.

## Supplementary materials

**Fig. A1.**
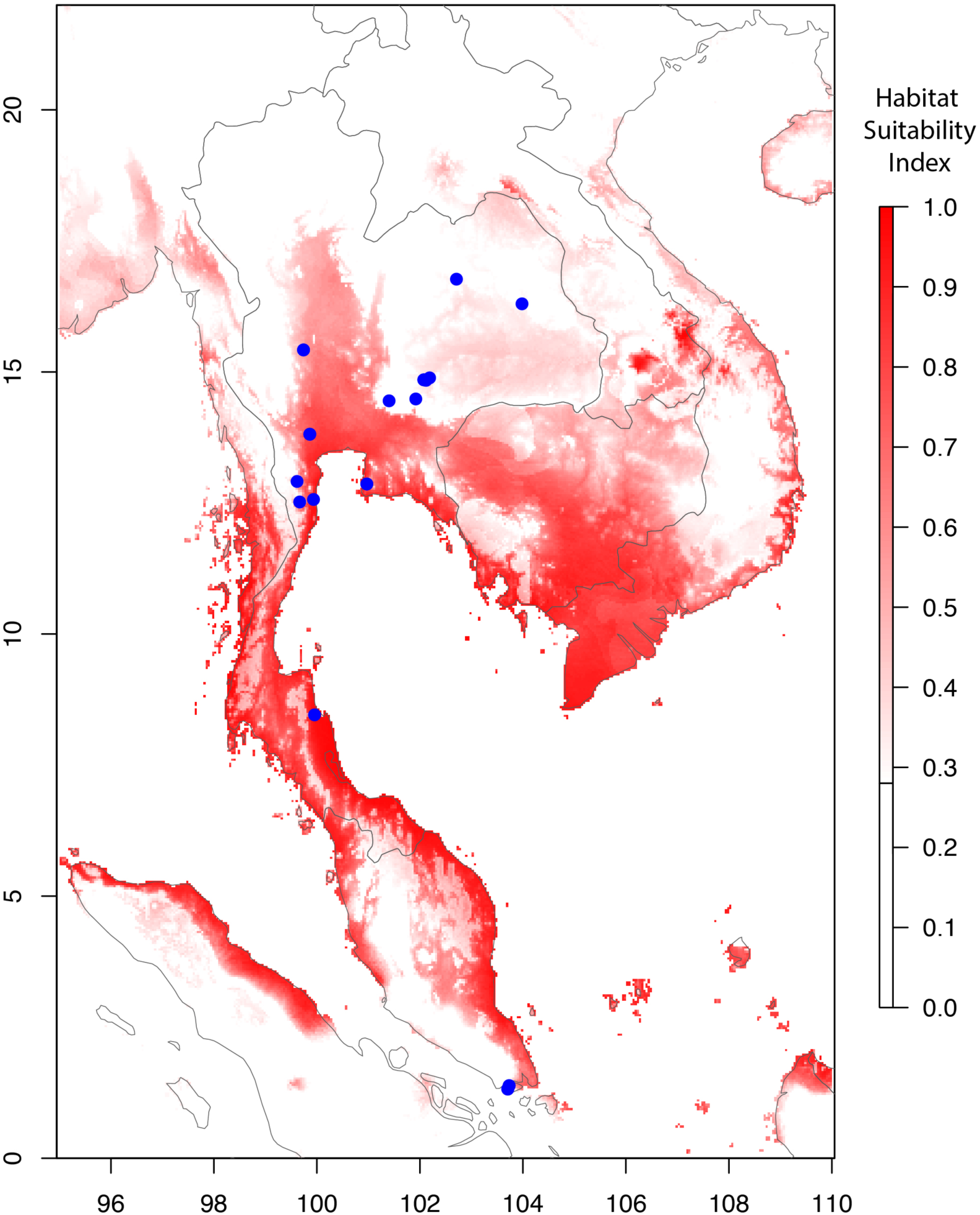
Species distribution model for *Iguana iguana* in Singapore, Thailand and surrounding countries for 2050 with the AC model and rcp45 for greenhouse gas emission (Fick and Hijmans 2017). Models used an absence threshold of 0.15 and habitat suitability ranges from high (dark red) to low (white). Blue dots represent locations with *I. iguana* presence.

**Fig. A2.**
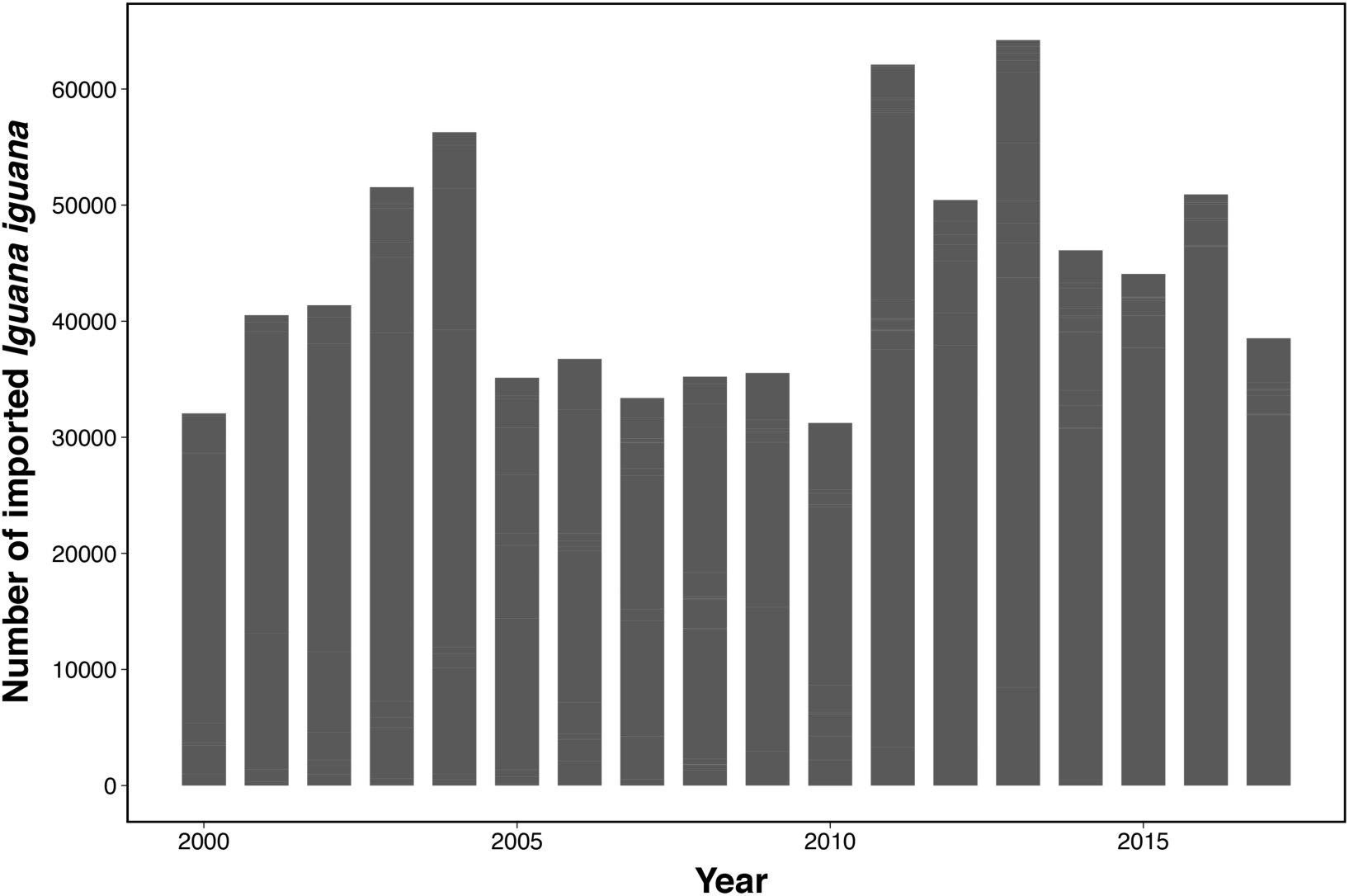
Reported CITES imports of *Iguana iguana* to Asian countries between 2000– 2017 (CITES Database 2019)

**Fig. A3.**
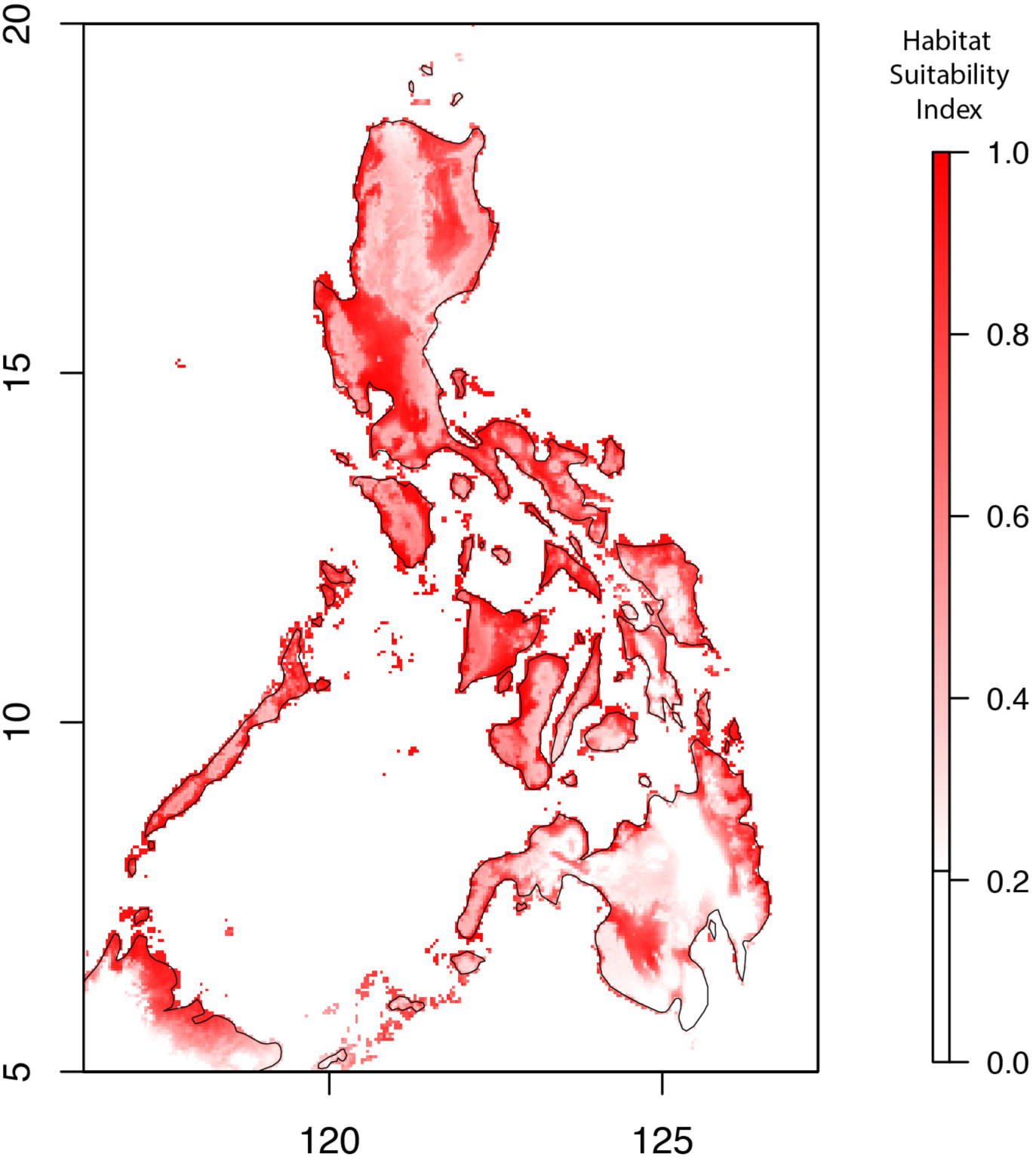
Species distribution model (as habitat suitability) for *Iguana iguana* in Philippines under current climate, with absence threshold 0.17. Habitat suitability ranges from high (dark red) to low (white).

**Table A1.**
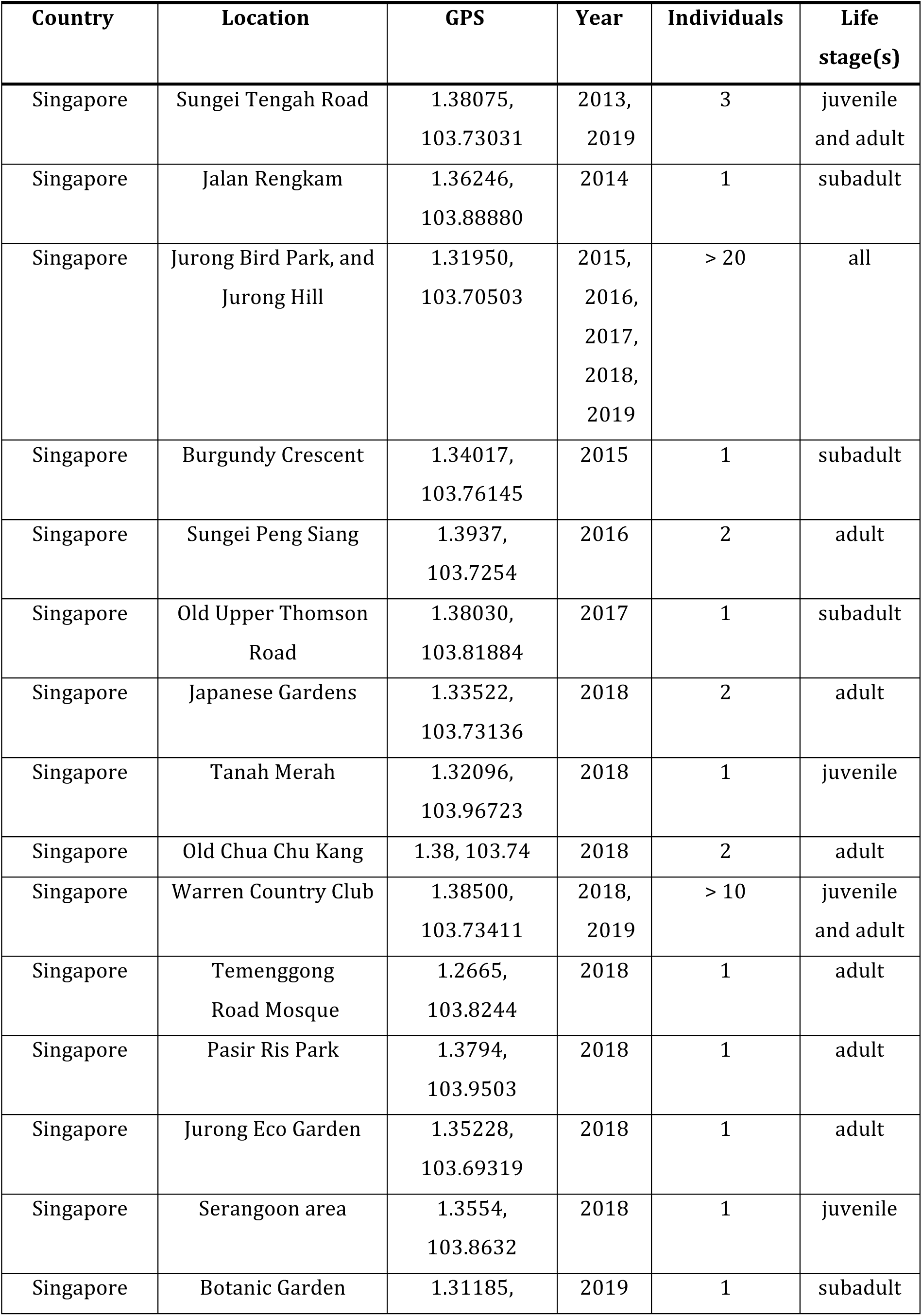
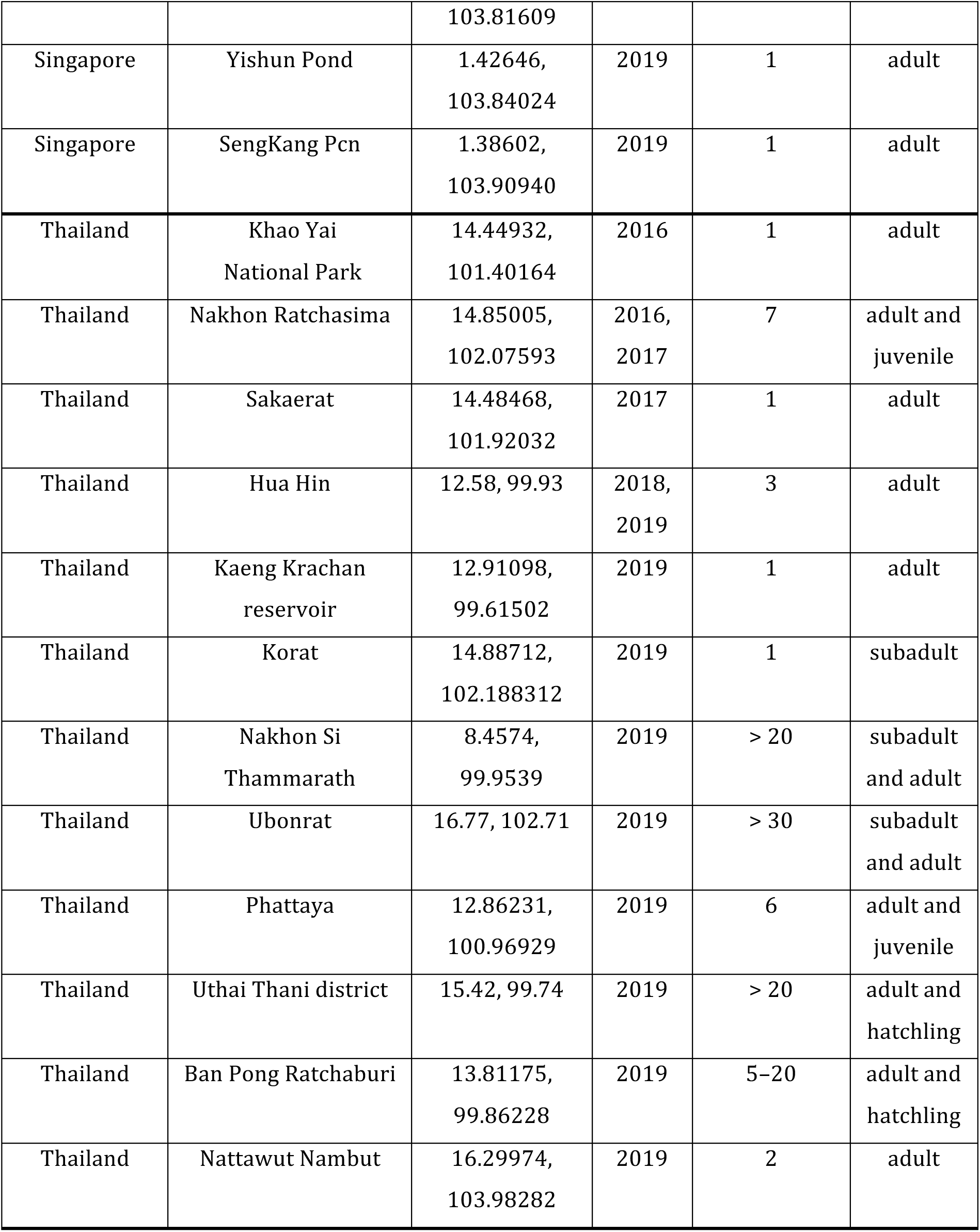
*Iguana iguana* sightings in Thailand and Singapore including location, year, number of individuals, and life stage(s) present

